# Multivariate Trait Evolution: Models for the Evolution of the Quantitative Genetic G-Matrix on Phylogenies

**DOI:** 10.1101/2024.10.26.620394

**Authors:** Simone P. Blomberg, Michelle Muniz, Mai Ngoc Bui, Cooper Janke

**Affiliations:** School of The Environment, University of Queensland, St. Lucia, 4072, Queensland, Australia; Institut für Applied and Computational Mathematics, Bergische Universität Wuppertal, Gaußstrasse 20, D-42119, Wuppertal, Germany; School of Business, British University, Ecopark, Hung Yen, Vietnam

**Keywords:** quantitative genetics, macroevolution, diffusions, Lie group theory, Riemannian geometry, Bayesian

## Abstract

Genetic covariance matrices (G-matrices) are a key focus for research and predictions from quantitative genetic evolutionary models of multiple traits. There is a consensus among quantitative geneticists that the G-matrix can evolve through deep time. Yet, quantitative genetic models for the evolution of the G-matrix are conspicuously lacking. In contrast, the field of macroevolution has several stochastic models for univariate traits evolving on phylogenies. However, despite much research into multivariate phylogenetic comparative methods, analytical models of how multivariate trait matrices might evolve on phylogenies have not been considered. Here we show how three analytical models for the evolution of matrices and multivariate traits on phylogenies, based on Lie group theory, Riemannian geometry and stochastic differential (diffusion) equations, can be combined to unify quantitative genetics and macroevolutionary theory in a coherent mathematical framework. The models provide a basis for understanding how G-matrices might evolve on phylogenies, and we show how to fit models to data *via* simulation using Approximate Bayesian Computation. Such models can be used to generate and test hypotheses about the evolution of genetic variances and covariances, together with the evolution of the traits themselves, and how these might vary across a phylogeny. This unification of macroevolutionary theory and quantitative genetics is an important advance in the study of phenotypes, allowing for the construction of a synthetic quantitative theory of the evolution of species and multivariate traits over deep time.

**Lay Summary:** We unite Quantitative Genetics, the major mathematical theory of multivariate quantitative trait microevolution, with the mathematical theory of multivariate macroevolution. To do this, we allow the key component of quantitative genetic theory, the matrix of additive genetic variances and covariances (the G-matrix) to evolve along evolutionary trees. This is an advance because the G-matrix is assumed to be constant in quantitative genetics (for convenience), but it has been recognised that it evolves on macroevolutionary timescales (in deep time). Uniting Quantitative Genetics with macroevolutionary theory allows for a more complete mathematical description of Darwin’s theory of evolution, and allows for further testing of evolutionary hypotheses.

## Introduction

The field of quantitative genetics is concerned with the evolution of traits over a small number of generations (McGuigan, 2006). A fundamental focus of research for quantitative genetics is the G-matrix, defined as the matrix of additive genetic variances and covariances of traits, usually derived from breeding experiments (Walsh and Lynch, 2018; Lynch and Walsh, 1998; Arnold, 2023). The G-matrix is an approximation of the underlying genetic architecture (Hansen, 2006). Exactly how and why it evolves are topics of current research (Arnold et al., 2008; Delahaie et al., 2017; McGlothlin et al., 2022). The matrix can be used to compute the predicted evolutionary response to selection of a collection of (multivariate) traits, using the multivariate breeders’ equation (Lande, 1979; Lande and Arnold, 1983):

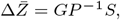

where 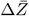 is a vector of responses to selection, with length *d*, the number of traits. *G* is the G-matrix, a *d × d* matrix of additive genetic variances and covariances. *P* is the phenotypic variance-covariance matrix, and *S* is a vector of “selection differentials” (Falconer and Mackay, 2009), also of length *d*. In a more compact form, the breeder’s equation can be formulated as:

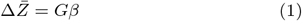

where *β* is a vector of partial regression coefficients relating traits to fitness (Lande and Arnold, 1983).

In Equation 1, G is taken to be constant over the timespan of a few generations; however Equation 1 is not appropriate for longer timespans where G likely evolves (Bégin and Roff, 2004; Steppan et al., 2002; Roff, 2000; Eroukhmanoff and Svensson, 2011; Arnold et al., 2008).

The field of macroevolution also deals with the evolution of traits, but over deep time, that is, over millions of years and across many speciation events. Quantitative macroevolutionary studies typically assume that evolution occurs according to some stochastic process, most often a diffusion (but see Landis et al., 2012; Bartoszek, 2017; Duchen et al., 2017; Landis and Schraiber, 2017). The stochastic differential equation (SDE) for an Itô diffusion process has the general form (Øksendal, 2010):

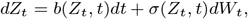

where *b*(*·*) and *σ*(*·*) are the “drift” and “diffusion” functions of a (univariate) trait *Z* at time *t. W*_*t*_ is a Wiener process, that is, Brownian motion (BM) at time *t*. This formulation is very general and many diffusion models have been proposed for macroevolutionary problems (Blomberg et al., 2020). Univariate models have been the focus of most macroevolutionary research, with some exceptions (Butler and King, 2004; Arnold et al., 2008; Bartoszek et al., 2012; Adams and Collyer, 2019; Clavel et al., 2015). Pure Brownian motion is the simplest and most used diffusion model in macroevolutionary studies, though BM should be treated carefully in the multivariate case. See Mörters and Peres (2010) for a discussion of the differing behaviour of multivariate BM processes.

### Lie Group Theory

To allow G-matrices to evolve on phylogenies, we need to make sure that the changes that occur during evolution of G in our models occur such that the properties of G remain unchanged. That is, we need to ensure that all G-matrices are symmetric and positive definite. A naive approach would be to allow the elements of G to evolve according to their own process but that would mean that G would very quickly become asymmetric and non-positive definite; the matrices could no longer be interpreted as covariance matrices.

We use Lie group theory to enforce the restriction that covariance matrices, which are symmetric and positive-definite, remain so during our simulation and data analysis experiments. (Belinfante et al., 1966; Iserles et al., 2000; Stillwell, 2008). We particularly recommend Stillwell (2008) as an introduction to Lie theory for non-specialists. Lie group theory is well-known in physics, where it is used to study symmetry relations in physical theories (Das and Okubo, 2014). However, there have been few applications of Lie group theory in biology (but see Fernandez-Sánchez et al., 2015; House, 2012; Shang, 2012; Sumner et al., 2012; Shang, 2013).

Recent advances in the application of Lie group theory to the evolution of covariance matrices according to SDEs show that in order to evolve matrices with the above properties (symmetric, positive-definite), it is much easier to evolve matrices on the corresponding Lie algebra, which is the tangent space to the matrix Lie group at the identity (Muniz et al., 2022b,a, 2023). Matrices can be transferred onto the Lie algebra using the matrix logarithmic map (Bui, 2022; Bui et al., 2023). While the set of covariance matrices is not closed under matrix multiplication or addition, the space of covariance matrices does form a group if they are endowed with the appropriate metric, either the Log-Euclidean metric or the Fisher-Rao Affine Invariant metric (Arsigny et al., 2007; Bui et al., 2023). If the Log-Euclidean metric is used, then the space of covariance matrices is a Lie group.

The results on this tangent space can then be transferred back to the associated Riemannian manifold (Petersen, 2006; Bhatia, 2015) using the matrix exponential map 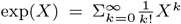 (Hall, 2015; Bui, 2022; Bui et al., 2023). Hence, we can exploit the relationship between the manifold and the tangent space in order to build matrix-evolutionary models.

### An Isospectral Model of Evolution

Without loss of generality, consider two traits evolving according to their own SDE where *b*_1_(*·*), *σ*_1_(*·*) and *b*_2_(*·*), *σ*_2_(*·*) are the drift and diffusion functions for trait 1 and trait 2, respectively. A system of *correlated* SDEs can be expressed as:

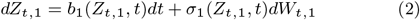

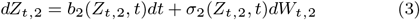

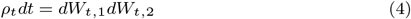

where *ρ*_*t*_ is the additive genetic correlation between Wiener processes *W*_1_ and *W*_2_ at time *t*. Equation 4 can be derived using basic stochastic calculus and Itô’s formula (e.g. Wiersema, 2008, p. 77). Parameter *ρ*_*t*_ can be derived from the G-matrix as

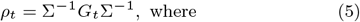

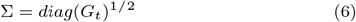

Each trait *Z*_*t*_ evolves according to its own (univariate) diffusion with its associated Brownian motion. It is these Brownian motions, one for each trait, that are correlated according to Equation 5, not the entire diffusion. So our models are not simply multivariate diffusion equations. Each trait has its own univariate diffusion, and *G* (via *ρ*) enters the model using equation 5. Our formulation allows each trait to have its own diffusion, which may not even be Gaussian (Blomberg et al., 2020). This is important because traits may be better modelled using non-Gaussian processes, but the relationship between the Brownian motions are easy to construct because Brownian motion itself *is* Gaussian.

We wish to include the evolution of *ρ*_*t*_ to study the evolution of the G-matrix at each time, *G*_*t*_. To do so, we must be careful that our evolutionary model ensures that, as mentioned above, like all covariance matrices G remains symmetric, real and positive- (semi)definite. This requires that all eigenvalues of G should be non-negative. In particular, if Ω_*t*_ is a skew-symmetric matrix solution to the following SDE:

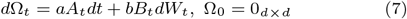

where *a* and *b* are constants to be estimated, *A*_*t*_ and *B*_*t*_ are time- dependent skew-symmetric matrices, and *W*_*t*_ is a scalar Wiener process, then

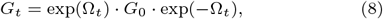

where **G**_0_ is the G-matrix at the root of the tree. Variable **G**_*t*_ will have the same (non-negative) eigenvalues as **G**_0_ by construction, and hence this is termed an “isospectral” model (Muniz et al., 2022a).

Equations 2 to 8 together form a model of multivariate trait evolution, with the G-matrix evolving on the matrix Lie group manifold. As well as the evolution of *G*, we can also simulate the vector of traits (*Z*), using univariate SDE models, one for each trait. Hence, we can model both G and the mean trait values across species. Since the traits follow their own evolutionary model constrained by the G-matrix, as the G-matrix evolves, the traits can be made to evolve along the phylogeny using consecutive updates of the G-matrix at each time step *t*.

This isospectral model can be used to evolve covariance matrices because the skew symmetric matrices (Ω_*t*_, constructed from the “mean” matrix for calibration) are closed under addition (“+”), and an Itô diffusion process is a series of additions, so long as the parameters themselves (Ω_*t*_) are skew-symmetric matrices. Since Ω_*t*_ is skew-symmetric, the matrix exp(Ω_*t*_) is in the special orthogonal group *SO*(*d*), which is a Lie group. Equation 8 is thus an action of the Lie group *SO*(*d*) on the convex cone of covariance matrices (J. Sumner pers. comm.).

Although the above model is isospectral, when using either the Log-Euclidean metric or the Fisher-Rao Affine Invariant metric, points on the manifold geodesics (ie covariance matrices) have their determinants resulting from the linear interpolation in the domain of the logarithm (Bui et al., 2023). This means our simulated covariance matrices need *not* be isospectral, and we can eliminate the need for skew-symmetric matrices as model parameters. Next, we introduce two non-isospectral models that are already popular for univariate traits, but which we extend to covariance matrices.

### An Ornstein-Uhlenbeck model for Covariance Matrices

One of the most popular univariate trait evolutionary models is the Ornstein-Uhlenbeck process (OU; Bartoszek et al., 2012; Hansen, 1997; Hansen and Martins, 1996) (but see Cooper et al., 2015;

Cornuault, 2022; Ho and Ané, 2014) which describes a process that is “mean-reverting:” as the process moves away from its central tendancy (*µ*), it gets “pulled” back towards the central tendency with a “force” *α*. Stochasticity is introduced with the addition of white noise to the system. Hence, the SDE for the OU process can be written as:

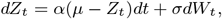

where *σ* is usually interpreted as the rate of evolution. Parameter *α* is interpreted as the strength of stabilising selection on trait *Z*, but if the process starts outside its (Gaussian) stationary distribution, directional selection can be observed as the process returns to equilibrium around *µ*. There are several univariate models that have OU-like reversion behaviour, and they have different stationary distributions, but we do not consider them here although we recognise that these models may prove to be valuable extensions (Blomberg et al., 2020).

The success of the OU process in modelling aspects of trait evolution on the adaptive landscape (Arnold et al., 2001; Svensson and Calsbeek, 2013; Bartoszek et al., 2012; Clavel et al., 2015), and the existence of models of covariance evolution as described here, suggest that it may be interesting to derive an OU-type model for covariance matrices. A matrix version of the OU process is:

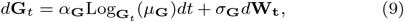

where **G**_*t*_ is the covariance matrix at time *t, α*_**G**_ is a *d*^*^ *× d*^*^ dimensional real matrix of reversion parameters, where 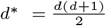, *µ*_**G**_ is the long-term mean covariance matrix, and *σ*_**G**_ is a positive-definite matrix of diffusion parameters, of size *d*^*^ *× d*^*^. **W**_*t*_ is a *d*-dimensional multivariate Brownian motion. 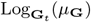is the logarithmic map from **G**_*t*_ evaluated at *µ*_**G**_. To implement this model, we use the same general approach as for the isospectral model. Instead of modelling on the covariance manifold (it is a convex cone; Hill and Waters, 1987), we model the evolution of the covariance matrix on the tangent space. For the isospectral model, we took the tangent of the manifold at the identity. However, the tangent does not have to be taken at the identity, and the starting values for the covariance matrix evolution can be mapped to the tangent space from any point on the manifold using the logarithmic map (defined as the inverse of the exponential map) with an appropriate metric (Bui, 2022; Bui et al., 2023). Here we use the Fisher-Rao Affine Invariant metric (Rao, 1945), where the distance *δ* between two covariance matrices P and Q on the manifold is:

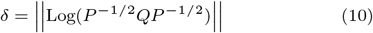

where ||*·*|| is the Frobenius matrix norm and Log is the matrix logarithm. Results can then be transferred back from the tangent space using the Fisher-Rao metric, the tangent point and the matrix exponential map.

### Brownian Motion of Covariance Matrices

Since the Brownian Motion model (BM) is just an OU model with *α* = 0, we can fit a matrix BM using the same computational methods as for the OU model. It has been pointed out that G-matrices cannot evolve by Brownian Motion (L. Revell, pers. comm., cited in McGlothlin et al., 2018). This is because covariance matrices are constrained to be positive-semidefinite, so if the elements of the matrix are allowed to evolve by BM in ordinary ℝ ^*d*^ then the matrices will “wander” off the manifold and the resulting matrices will not be positive-semidefinite (see Fig. 2 and Fig. 3 of Marjanovic et al. (2015) for a graphical representation of this phenomenon). This is avoided by allowing the log-mapped matrices to evolve on the tangent space, and transforming the resultant matrices back to the manifold of covariance matrices using the exponential map and the Fisher-Rao metric.

**Fig. 1:**
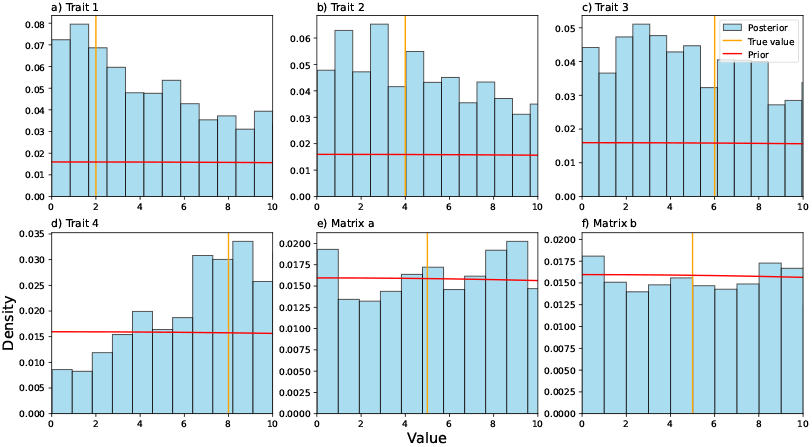
Prior distributions and simulated posteriors for simulated data, 4 traits and one 4 x 4 matrix on a tree with 50 leaves (a) to d)). The parameter being estimated is the *α* associated with the OU process for the traits, and parameters *a* and *b* for the Isospectral matrix model (Subfigures e) and f)). Orange lines represent the “true” value. The prior distributions over likely values of the parameters are represented by the red horizontal lines.

**Fig. 2:**
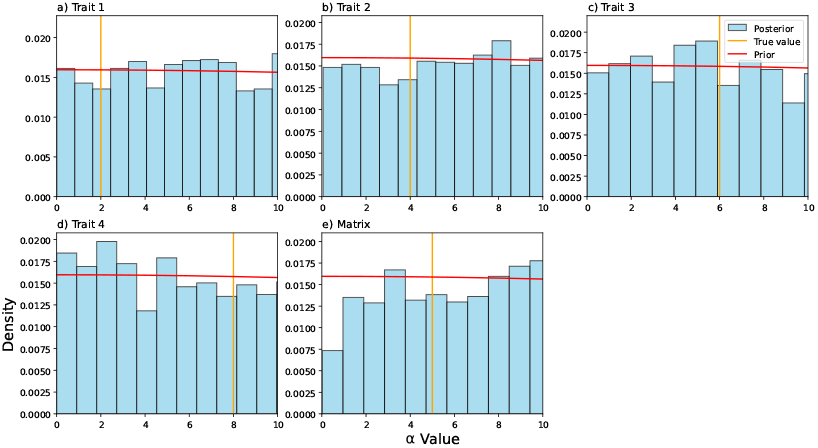
Prior distributions and simulated posteriors for simulated data, 4 traits and one 4 x 4 matrix on a tree with 50 leaves. The parameter being estimated is the *α* associated with the OU process. Orange lines represent the “true” value. The prior distributions over likely values of the parameters are represented by the red horizontal lines.

**Fig. 3:**
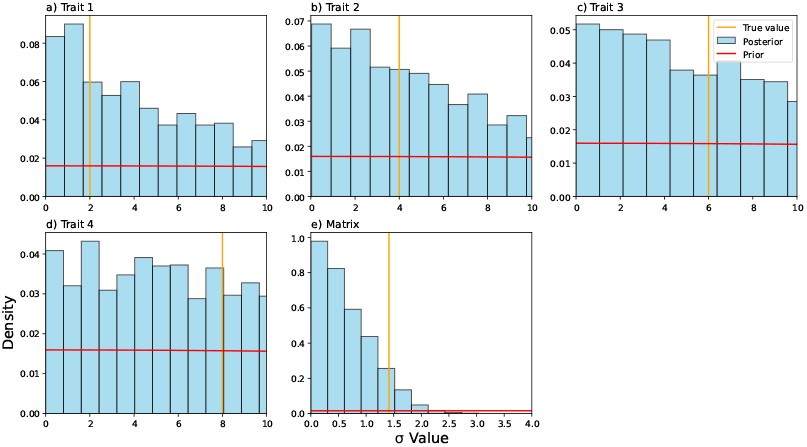
Prior distributions and simulated posteriors for simulated data, 4 traits and one 4 x 4 matrix on a tree with 50 leaves. The parameter being estimated is the *σ* associated with the Brownian motion. Orange lines represent the “true” value. The prior distributions over likely values of the parameters are represented by the red horizontal lines. Note especially subfigure d), where the x axis scale is greatly reduced and estimation is concentrated on very low values, markedly different from the prior.

**Fig. 4:**
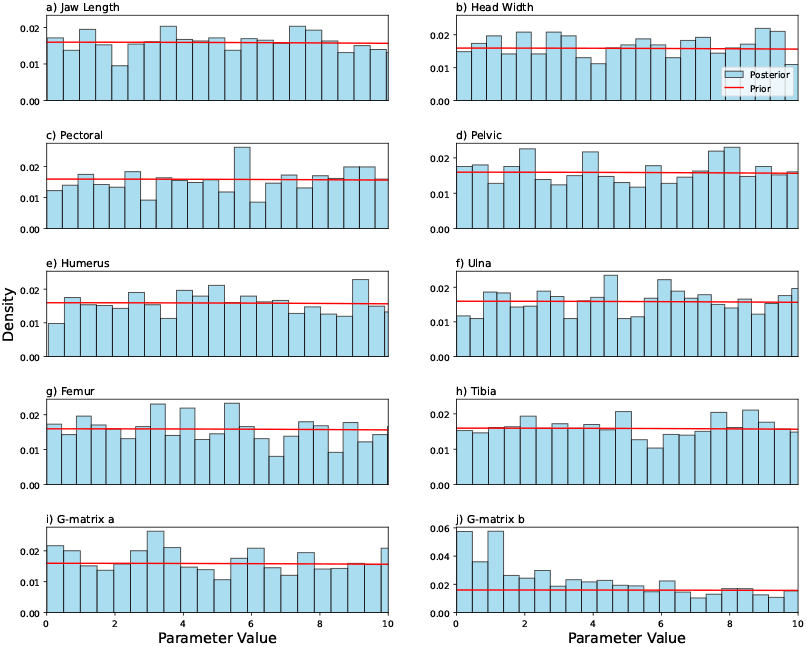
Prior and posterior distributions for the estimate of parameters using an OU model for traits and an ISO model for G-matrix evolution based on the *Anolis* dataset. This simulation allowed for a different *α* value for each trait. a-h) - distributions for the estimate of the trait’s *α* values based on the *Anolis* dataset. i-j) - distributions for the estimate of the matrix *a* and *b* parameters respectively.

### Data Requirements and Assumptions

The required data necessary for running the models are: a. A phylogeny for the species being examined, b. trait data (more than one trait), and c. the G-matrix of the traits under study, one matrix for each species. Additionally, data from fossils can be useful in improving inference for comparative analyses (Slater et al., 2012). We suspect that it is probably impossible to attain fossil G-matrices, however in some circumstances, the matrix of phenotypic covariances (P) can be used instead of G (Roff, 1995; Cheverud, 1988; Sodini et al., 2018). This substitution may be necessary if considering fossil taxa or non-model organisms. Our new models can be used on *any* covariance matrices. However we restrict our attention to the G-matrix as that is of the most theoretical importance.

The main assumptions of our models are that for all species, and traits *Z*, the G-matrix and the phylogeny are known without error. However, more complicated extensions can be easily envisaged, for example accounting for various model uncertainties by drawing samples from prior distributions of matrices, traits, or trees.

Of crucial importance, it is necessary to have a falsifiable scientific hypothesis about how and why parameters might hold the values that they do over the phylogeny (Popper, 1934). We wish to discourage “fishing expeditions” that search to optimise parameters across phylogenies without any reference to a particular scientific hypothesis. While such studies may be useful in generating hypotheses, these hypotheses cannot be tested with the data from which they are derived.

## Methods

We implemented the above models in the computer language, Julia (Bezanson et al., 2017). Julia is an ideal choice for analysing models of this type, as there are excellent facilities for solving differential equations (Rackauckas and Nie, 2017b), including SDEs (Rackauckas and Nie, 2017a), as well as packages for computations on manifolds (Congedo, 2024), linear algebra and using phylogenies (Reeve et al., 2023). Julia is fast, and is an open-source solution for technical and scientific computing. As a motivating example, we analysed the evolution of eight traits and G-matrices for seven species of *Anolis* lizards, using data from McGlothlin et al. (2022). We recognise that currently, datasets consisting of G-matrices for a group of species with a known phylogeny are extremely rare (We could only find the *Anolis* example). We hope that future studies will become more plentiful, with a larger sample of species. Previous work on phylogenetic signal suggests that a sample size of approximately 20 species is necessary to detect phylogenetic signal if it is present (Blomberg et al., 2003). It is unlikely that our seven *Anolis* species will provide enough power to estimate parameters with any precision. We used simulation to test our models’ ability to recover “true” parameter values, using a simulated phylogeny with 50 tips (leaves) and four traits.

For many SDE models, transition densities and thus model likelihood functions are either intractable or unknown. For this reason, researchers have generally resorted to approximations of the likelihood, e.g. the Euler approximation (Iacus, 2008; Blomberg et al., 2020), although if the univariate OU process is considered, the transition density (and hence the likelihood) is Gaussian and the usual maximum likelihood and Bayesian approaches can be used. Another approach, taken here, is to give up on likelihood inference altogether and use Approximate Bayesian Computation (ABC; Beaumont, 2010; Csilléry et al., 2010; Lintusaari et al., 2017; Toni et al., 2009). We use ABC here, even though we are demonstrating the theory that for the OU model is a system of multivariate OU processes for which the transition density is known. This is because we wish to keep the analytical methods general so more complicated models with unknown transition densities may be used in the future (Blomberg et al., 2020).In fact, while the univariate OU model can be analysed using maximum likelihood, the matrix models cannot, due to the fact that for the matrices, evolution is not in *R*^*d*^ but on a manifold. A simple ABC rejection algorithm involves four steps: 1.

Draw parameter values from a prior distribution. 2. Use these parameters to simulate data. 3. Compare the simulated data to the real data. 4. If the simulated and real data are sufficiently similar, the current parameter values (particles) are accepted into the posterior distributions for the parameters. This process is repeated a large number of times, to build up posterior distributions for all the model parameters. The approach has had some success in constructing phylogenetic comparative methods (e.g. Bartoszek and Li`o, 2019; Beaumont, 2010; Haba and Kutsukake, 2019; Jhwueng, 2020). For our analyses, we used a Sequential Monte Carlo ABC approach (Sisson et al., 2007, 2009) based on the Sampling-Importance-Resampling (SIR) algorithm due to Rubin (1987, 1988). The SIR algorithm proceeds by recognising that the rejection algorithm is wasteful, since many simulation runs are discarded and do not contribute to the posterior distribution at all. The SIR algorithm makes use of all the simulations, but weights each particle according to its “importance,” which is a kernel density evaluated at the “distance” between the observed and simulated data values. We used a Gaussian kernel with *σ* = 3. The particles were resampled, with particle weights equal to their (normalised) importance, thereby producing an approximation to the posterior distribution. We used bespoke Julia code to implement ABC-SIR for our matrix simulation models. Our code is available on GitHub.

Central to the application of ABC to any problem is the question of how to measure the “similarity” of simulated data to observed data. Ideally, summary statistics for the comparison of “observed” to “simulated” data should be functions of sufficient statistics (Casella and Berger, 2024). In the present unusual case, the data themselves (trait values and covariance matrices) are a sufficient statistic, and we can compare the simulated trait values directly to the observed trait values using the difference between the simulated trait data and the observed trait data for each species (leaf).

There are many ways to describe the similarity of matrices (reviewed in Steppan et al., 2002). Here we use the Fisher-Rao Affine Invariant metric of the distance between two covariance matrices on the manifold (Equation 10). We recognise that other metrics could be useful for determining the similarities of simulated matrix and trait data to observed data (e.g. Phillips and Arnold, 1999). Some methods may be able to better describe *why* matrices differ (Aguirre et al., 2014). However we chose to keep the analysis relatively simple and as a demonstration, rather than provide more precise measures of matrix similarity. For our present purposes, the Fisher-Rao Affine Invariant metric has strong theoretical support (Arsigny et al., 2007; Bui et al., 2023).

Importance weights were calculated for both matrices and traits and then summed across all traits, matrices and species, to give one importance value for each proposed trait value (particle).

### Parameter Estimation

For both the simulation study and the *Anolis* data, initial trait values at the root were estimated as the species mean for each trait. For the Anolis study, the value of the G-matrix at the root was set to be the “ancestral” G-matrix, as per McGlothlin et al. (2022). These root values (traits and G-matrix) were regarded as fixed. For each model, we ran our ABC-SIR analysis 5000 times and resampled 10 000 times. For both the simulation study and the anoles, all *σ* values were set to 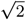for the OU model, where *α* was estimated. When estimating *α, σ* parameters were set to unity. When estimating the parameters from the isospectral model, we set all *σ* parameters to unity and fitted univariate OU models to the trait data.

We tried to use weakly informative priors wherever possible, to regularise the analytical problem and exclude the sampling of unreasonable parameter values (Lemoine, 2019). After considerable experimentation, a Truncated Half Normal distribution with *σ* = 50 was used for all parameters, which was essentially flat over the region of likely parameter values (less than 10 units).

For the *Anolis* study, we calibrated the skew-symmetric matrices for the isospectral model (the *A* and *B* parameters in using the “ancestral” G-matrix, derived in McGlothlin et al. (2022).This ancestral G was converted to a skew-symmetric matrix by setting the diagonal elements to zero and multiplying the upper-triangular elements by negative one. Time dependence was introduced by multiplying each matrix by *t* (time). We also set *A* = *B*. We used the same procedure for the simulated data study. In that case we used the traits and G-matrix as described in the Supplementary Material.

We scaled the phylogeny to have a height of one. For the tree topology, see Zheng and Wiens (2016) and McGlothlin et al. (2022). In all analyses, we used a step size of 0.01 time units along the tree.

For the OU model, we restricted the number of parameters to just one *α*_*G*_ for the G-matrix estimation, and one *α* for each for the trait estimations, to improve model identifiability. Similarly, for the isospectral model we used one *α* value for OU univariate trait models and we estimated the *a* and *b* parameters of the matrix isospectral model.

### Model Selection

We performed Bayesian model selection for each of the three models using the same model specifications and simulations as for the parameter estimation. The relative posterior probability for each model was approximated by the sum of the importances obtained for each model, before normalization. The model selection criterion was that the better model(s) have a higher total importance. The prior used for each model was a discrete Uniform distribution of span 3, since there were three models in the comparison set. Data were simulated under an OU model for the traits and the G-matrix. Our prediction was that OU would be preferred by our model selection criterion.

## Results

### Simulations

#### Parameter Estimation

We simulated multivariate data (4 traits) along an artificial tree with 50 leaves, under the isospectral model, BM and OU models for both the traits and their G-matrix (figs. 1 to 3). The effectiveness of parameter estimation (For BM, *σ* and for OU, *α*) was variable but in most cases the prior distributions moved towards the true values such that the posterior distributions easily covered the true values and there was a marked concentration of posterior density around the true values. The exception to this was where smaller values of the matrix *α* were strongly preferred (Fig. 2 e)). Further experiments (not shown) suggested that parameter estimation is easier for *α* if the starting trait value at the root is in the tails of the stationary distribution, indicating that the inclusion of fossil information is useful, particularly when fossils look unlike modern forms (Benton, 2015; Slater et al., 2012).

#### Model Selection

For the simulated data, ABC-SIR model selection chose the correct model (OU; Table 1).

**Table 1.** Model selection results comparing OU, BM and the isospectral model for data simulated under an OU model of evolution. Values are the relative percentages of the importances across the three models. As expected, The OU model was roundly favoured (having higher total importance), and BM was least supported. See Discussion.

| OU | BM | ISO |
| --- | --- | --- |
| 65.38 | 13.54 | 21.08 |

### Anolis Analysis

#### Parameter Estimation

The estimates of the trait *α* parameters were not precise, and not very different from the prior distributions (Figs. 5 - 6). This is to be expected as the sample size (seven species) was very small and there is little information about the dynamics (ie. *α*) in data based only on extant species (Blomberg et al., 2020). There were some notable exceptions. Concentration of the posterior distribution at small values for the isospectral *b* parameter, support a conclusion that *b* is small in this dataset. For the OU analysis, there are no strong departures from the prior distributions for *α* for the traits, but there is strong support for small values for the matrix *α* (Fig. 5). For the BM analysis, there was a strong concentration of the posterior density at small *σ* values for the matrix parameter (Fig. 6)

**Fig. 5:**
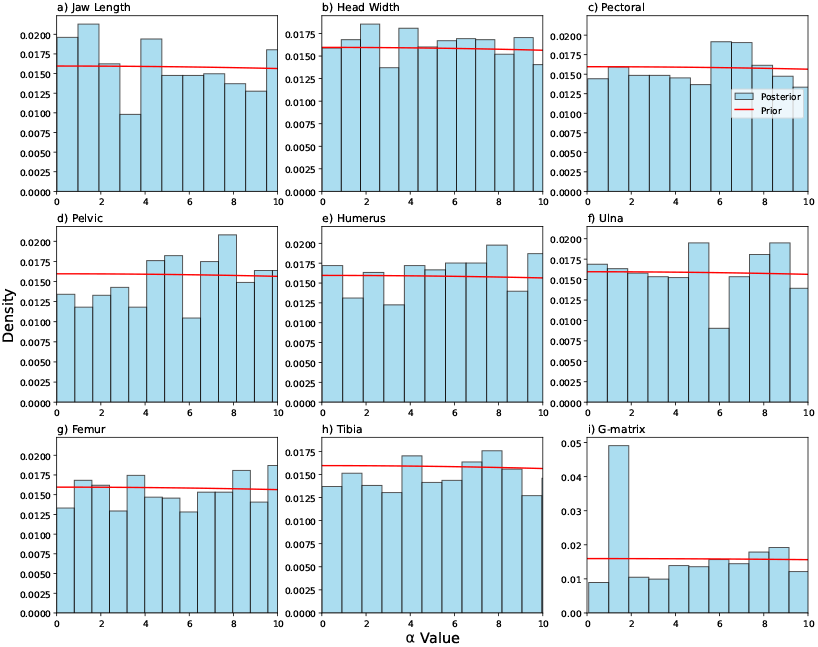
Prior and posterior distributions for the estimate of parameters using an OU model for trait and G-matrix evolution based on the *Anolis* dataset. This simulation allowed for a different *α* value for each trait. a-h) - distributions for the estimate of the trait’s *α* values based on the *Anolis* dataset. i) - distributions for the estimate of the matrix *α* based on the *Anolis* dataset.

**Fig. 6:**
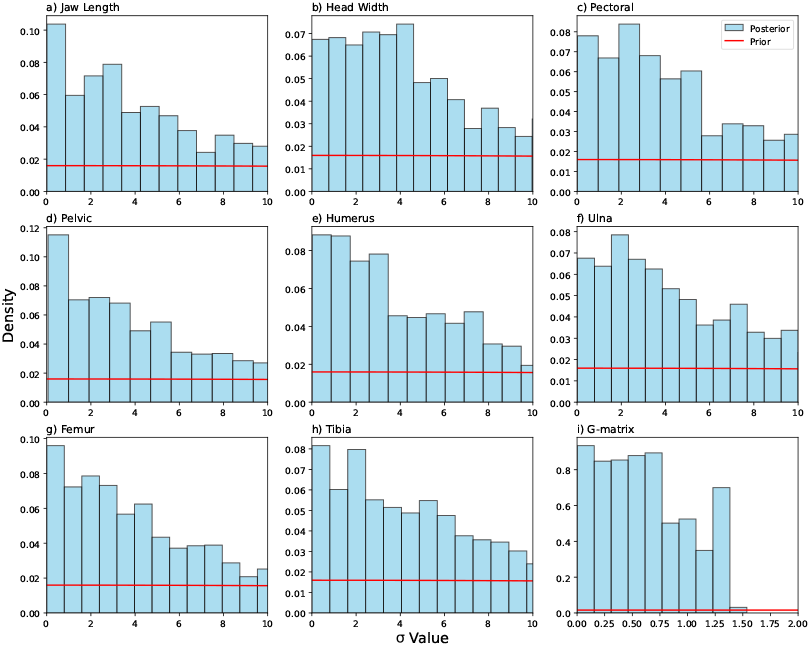
Prior and posterior distributions for the estimate of parameters using a BM model for traits and a BM model for G-matrix evolution based on the *Anolis* dataset. This simulation allowed for a different *α* value for each trait. Note especially the very large concentration around small values in subfigure i).

Given the lack of biological realism (using a single *α* for the G-matrix. We do not expect each element of the G-matrix to have the same *α*), we view this result as a demonstration of what is possible in this model-fitting framework, rather than a definitive answer to the question of the strength of stabilising selection on the G-matrix. We do note that such a question could not be answered except by using a quantitative macroevolutionary model for the G-matrix.

#### Model Selection

Model selection for the *Anolis* data clearly favoured the isospectral model over OU and BM (Table 2). Support for OU was very weak.

**Table 2.**
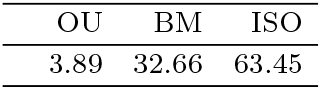
Model selection results comparing OU, BM and the isospectral model for 7 *Anolis* species. Values are the relative percentages of the importances across the three models. The isospectral model was roundly favoured, and OU was least supported. See Discussion.

## Discussion

We have shown how to effectively model the macroevolution of traits over deep time, including the information on covariance matrices (the G-matrix) to determine the correlation among traits. Further, we have been able to model the evolution of the G-matrix itself. This fills an important gap in the macroevolutionary theory for multivariate traits, as the G-matrix probably does evolve over time, as evidenced by the fact that G-matrices can be different for related taxa (e.g. the *Anolis* case, McGlothlin et al., 2018, 2022). Central to the mathematical description of the evolution of G-matrices is our use of Lie group theory and Riemannian geometry to ensure that the evolution of G results in matrices that are symmetric, positive-definite and are thus interpretable as covariance matrices. We have developed three models that can represent the evolution of covariance matrices, two based on the univariate case, BM and OU, and the isospectral model which preserves the eigenvalues of the ancestral covariance matrix. BM and the isospectral model may both serve as null models in further studies of the evolution of G-matrices on phylogenies. OU may be particularly useful in testing for selection on G-matrices, and to determine how much of G-matrix evolution is constrained, by studying the *α*_**G**_ parameter.

Overall, we found these models difficult to fit to both simulated and real data (figs. 1 to 6). While the posterior densities of simulated model parameters usually included the true value, the densities were often wide and not very much different from their priors. Subsequent computer experiments (not shown) starting simulations from root values outside of the OU stationary distribution, which may be interpreted as an initial burst of directional selection followed by stabilising selection, appeared more promising for the estimation of the OU parameter *α*. Hence, it may be possible to obtain better model fits if the dataset includes traits of fossil taxa as well as extant taxa, and if optima differ in different places in the phylogeny, or if the root values are very different from the tip species values. Of course, this will probably necessitate using phenotypic covariance matrices rather than G-matrices.

In the model selection procedure, the OU model was preferred over the others, in the case where data were generated under an OU model (Table 1). The isospectral model was strongly preferred in the *Anolis* example (Table 2). However, the model selection results are best interpreted along with the parameter estimates, and given the lack of precision in those, we think the model selection results should be treated with caution.

The relationship between macroevolution and microevolution has long been a topic of speculation (Serrelli and Gontier, 2015), with some researchers convinced that macroevolution is in some fundamental way different from microevolution (e.g. Gould, 1985). Alternatively, others claim that macroevolution is just microevolution “writ large” (Hendry and Kinnison, 2001; Reznick and Ricklefs, 2009). It is clear now that statistical genetics and phylogenetics have very deep common roots (Schraiber et al., 2024). The time is ripe for a synthetic mathematical theory that unites macroevolutionary theory in terms of stochastic processes of trait evolution acting cross many generations and speciation events, and microevolutionary theory, the theory of phenotypic evolution on small timescales, in the form of evolutionary quantitative genetics (Rolland et al., 2023). The models proposed here are a step in that direction. These models could be extended in various ways, e.g. the model parameters might be allowed to vary over time, producing models that allow for different selection regimes (*α*), different optimal trait values (*µ*) different values of *σ* (Beaulieu et al., 2012; Ingram and Mahler, 2013; Beaulieu and O’Meara, 2023; Revell, 2021), the use of non-Gaussian OU-like models (Blomberg et al., 2020), the application of “early burst” models of adaptive radiations (Gavrilets and Losos, 2009), or even the importance of phenotypic plasticity in macroevolution (de Souza Silva et al., 2023). Diffusion models provide a rich framework for studying macroevolution, and there are many possibilities open to answer questions using a wide variety of models. The models presented here can form the basis for further studies. This new modelling paradigm will allow the testing of new hypotheses of trait variation, including hypotheses about the evolution of phenotypic integration (Pigliucci, 2003; Eble et al., 2004; Caetano and Harmon, 2017) and the coevolution of the strength of the association among groups of traits (Steppan, 2004).

### Biological Interpretations of Theory

For each of the three models presented here, one can ask the question, “what do they mean?” That is, what are the biological interpretations of the parameters? The coupled univariate trait models (equations 2 and 3) have the same interpretation as in the usual univariate uncoupled case. *σ* is the “rate” of evolution, that is, Gaussian “noise” is introduced into the system by some scaling factor *σ*. This noise is the result of random mutation and genetic drift on the relevant genes that contribute to the phenotype, independent of selection. *σ* can also be interpreted as the scaling of the phenotypic response to strong selection, but in a randomly varying environment (See: Hansen and Martins, 1996). Both of these interpretations are likely to be somewhat incorrect, but if we wish to use diffusion models as models of evolution (which is the common practice), we are stuck with these interpretations. For the OU model, we have two extra parameters: *α* and *µ. α* is usually interpreted as the strength of stabilising selection around the central tendency (*µ*). This is a vast simplification of the underlying ecological processes that contribute to the multiple sources of natural selection that occur in populations. In essence, the ecology of a species that is relevant to selection is represented by a single number. We encourage macroevolutionary modellers to devise new ways of incorporating more realistic representations of selection that do better justice to the complexities of organisms’ ecology.

The interpretation of the matrix evolutionary models (equations 7 and 9) is more subtle. Equation 7 has two parameters, *a* and *b*, which are estimated. We cannot see a direct connection for these parameters to biology as can be seen in the univariate BM and OU models. However, we can consider what it means, biologically, to have isospectral G-matrices. If two matrices are isospectral ie. have the same eigenvalues, but different elements, then the differences must be evident in the matrices having different eigenvectors. The eigenvectors represent the genetic “lines of least resistance” to evolution through phenotypic space where natural selection can produce the greatest phenotypic response (Schluter, 1996; McGuigan et al., 2005). Hence, our isospectral model allows for the direction of likely evolution to change, but the magnitude of the genetic variation in the direction of this varying eigenvector remains constant. We think this is an interesting idea because it implies that lines of least resistance may alter independently of standing genetic variation, perhaps due to changing environments (Wood and Brodie III, 2015). This decoupling of eigenvalues and eigenvectors provides an interesting line of future research, perhaps using our isospectral model as a building block. It would be interesting to know whether this behaviour is common, and what could drive it.

In contrast, the OU and BM models of matrix evolution are more prosaic. The OU model uses a matrix of mean-reverting terms (*α*_**G**_), one for each unique element of the G-matrix, a “reversion” matrix (*µ*_**G**_) which acts as a central tendency to which the stochastic process returns, and *σ*_**G**_, a matrix of evolutionary rates. There is also a matrix of Brownian motions, one rate for each unique element in the G-matrix (*dW*_**t**_). The interpretations of the elements of *α*_**G**_, *µ*_**G**_ and *σ*_**G**_ are the same as for the univariate case. The only proviso is that here the matrix diffusion processes evolve on the tangent space to a Riemannian manifold. Hence, projecting the simulated matrices back onto the manifold results in trajectories on the manifold that are *not* equivalent to simple BM and OU processes on the matrix elements. They cannot be, as we are enforcing the restriction that the simulated matrices remain as covariance matrices.

We emphasise that there is no biological interpretation of the manifolds and tangent spaces. These spaces have no ontological existence and are present in the theory because of the necessity to evolve covariance matrices such that the new, evolved, matrices are also covariance matrices. In that sense, they are no more than a useful mathematical trick. Covariance matrices play an important role in the construction of modern evolutionary theory. This point is emphasised because we can now treat covariance matrices as traits in themselves, which means they are open to mathematical and biological study in the context of macroevolution.

## Conclusion

We suggest that there is another useful application of these stochastic process models: They provide us with a way to think about quantitative trait evolution. Even if these models prove difficult to fit to trait data, which is often the case (Cooper et al., 2015; Cornuault, 2022; Ho and Ané, 2014), they nevertheless have heuristic value. They allow us to draw a picture of the evolution of traits that is elegant, instructive, rigorous, and is a concise mathematical treatment of the known subject matter. In this sense, the statistical approach to trait evolution offered here is similar to the use of the Wright-Fisher model in population genetics (Fisher, 1930; Wright, 1931). It is unlikely that the Wright-Fisher model (and its extensions) is “true” in every respect (or at all), but nevertheless the heuristic value of the model in explaining the influences of drift, migration, mutation and selection of alleles in populations is profound. While our models are necessarily wrong, we believe they may be useful (Box, 1976). We think that statistical models can have a value of their own, beyond the usual role as serving as an adjunct to the testing of hypotheses about parameters (Sokal, 1965). They can reveal the processes of trait evolution.

## Data and code availability

All original Julia code is available at: https://github.com/simone66b/JMMenura. The *Anolis* data come from McGlothlin et al. (2022).

## Author contributions

S. P. B conceived the study, wrote code, and prepared the manuscript. M. M. and M. N. B. contributed to theory and approved the manuscript. C. J. wrote code, ran analyses and contributed to writing the manuscript.

## Acknowledgments

We thank D. Fisher, A. Hayes, B. Martin, D. Ortiz-Barrientos and M. Phillips for comments on a previous draft of the manuscript.

J. Sumner provided enlightening discussions on the intricacies of Lie group theory. D. B. Rubin suggested the ABC-SIR approach.

J. Schraiber, J. McGlothlin and one anonymous reviewer provided insightful comments and suggestions that markedly improved the manuscript.

