## Supplementary material for "Multivariate Trait Evolution: Models for the Evolution of the Quantitative Genetic G-Matrix on Phylogenies": Suplementary Tables

### Supplementary Material for Blomberg et. al. 2025. Evol. Letters.

| | Trait | | | |
|---|---|---|---|---|
| | 1 | 2 | 3 | 4 |
| 1 | 133.65 | 9.24 | -6.53 | -14.03 |
| 2 | 9.24 | 102.97 | -15.70 | 6.70 |
| 3 | -6.53 | -15.70 | 89.76 | 13.94 |
| 4 | -14.03 | 6.70 | 13.94 | 97.15 |

Table 1: Root G-matrix ($G_0$) for four traits for the simulated data analysis

| | Trait | | | |
|---|---|---|---|---|
| | 1 | 2 | 3 | 4 |
| 1 | 130.10 | -6.13 | -4.70 | -3.12 |
| 2 | -6.13 | 99.03 | 9.63 | -6.44 |
| 3 | -4.70 | 9.63 | 106.70 | 6.16 |
| 4 | -3.12 | -6.44 | 6.16 | 94.62 |

Table 2: $\mu_G$ for the four traits in the OU model for the simulated data analysis

| Trait | | | |
|---|---|---|---|
| 1 | 2 | 3 | 4 |
| 2.12 | 1.50 | 1.22 | 1.06 |

Table 3: Starting trait values ($Z_0$) for the simulated data analysis

| | Trait 1 | Trait 2 | Trait 3 | Trait 4 |
|---|---|---|---|---|
| 1 | 1.88 | 1.85 | 0.41 | 9.90 |
| 2 | 1.86 | -0.02 | 3.57 | 3.99 |
| 3 | 4.55 | -2.88 | 7.02 | 12.40 |
| 4 | 2.95 | 1.26 | 2.73 | 12.20 |
| 5 | 1.97 | 2.75 | 0.31 | 9.52 |
| 6 | 1.77 | 2.00 | 1.80 | 6.92 |
| 7 | 1.24 | 2.28 | 1.48 | 4.62 |
| 8 | 2.10 | -1.73 | 7.32 | 12.90 |
| 9 | 2.57 | 4.30 | -1.15 | 8.51 |
| 10 | 3.76 | 1.82 | 2.01 | 14.30 |
| 11 | 3.14 | 1.16 | 2.09 | 12.60 |
| 12 | 3.68 | 1.20 | 1.42 | 15.20 |
| 13 | 1.91 | -2.78 | 8.72 | 7.04 |
| 14 | 2.77 | -2.41 | 6.90 | 5.27 |
| 15 | 3.15 | -3.58 | 7.76 | 7.69 |
| 16 | 1.38 | -2.64 | 6.50 | 9.77 |
| 17 | 3.54 | -2.27 | 5.84 | 7.19 |
| 18 | 0.49 | -5.31 | 12.70 | 12.70 |
| 19 | -0.49 | 2.64 | 4.03 | 2.04 |
| 20 | 4.73 | -2.56 | 7.02 | 12.90 |
| 21 | 3.24 | -2.10 | 7.76 | 12.60 |
| 22 | 3.40 | -2.01 | 9.03 | 14.90 |
| 23 | 3.05 | -1.19 | 4.87 | 6.68 |
| 24 | 3.58 | -0.90 | 4.81 | 7.49 |
| 25 | 1.72 | 1.89 | 0.73 | 0.68 |
| 26 | 1.54 | 2.08 | -0.40 | -1.28 |
| 27 | -0.12 | -3.85 | 8.28 | 17.80 |
| 28 | -0.45 | -1.27 | 0.53 | 8.62 |
| 29 | -0.95 | 0.07 | -0.38 | 8.65 |
| 30 | -1.53 | -3.44 | 3.95 | 14.20 |
| 31 | -0.93 | 4.34 | -2.28 | -0.35 |
| 32 | -0.38 | 5.55 | -4.54 | -2.93 |
| 33 | -1.05 | 6.43 | -8.85 | -7.61 |
| 34 | -1.57 | 7.62 | -9.11 | -9.47 |
| 35 | -0.68 | 6.03 | -7.23 | -4.05 |
| 36 | -0.78 | 6.56 | -7.52 | -4.56 |
| 37 | -0.64 | 3.10 | -3.17 | 2.08 |
| 38 | 1.98 | 2.28 | 0.22 | 4.03 |
| 39 | 1.49 | 3.35 | -4.83 | 7.65 |
| 40 | 1.71 | 3.62 | -2.99 | 9.11 |
| 41 | 0.92 | 5.46 | -3.20 | 5.99 |
| 42 | 0.83 | 5.35 | -3.71 | 5.88 |
| 43 | 1.23 | 4.51 | -2.32 | 3.68 |
| 44 | 0.33 | 3.57 | -1.97 | 1.91 |
| 45 | -0.39 | 0.12 | -1.28 | 2.30 |
| 46 | 1.34 | 0.82 | 0.21 | 6.77 |
| 47 | -1.14 | 0.16 | -2.77 | 3.70 |
| 48 | 1.08 | 0.21 | -0.19 | 6.15 |
| 49 | -0.39 | 2.44 | -5.48 | 6.27 |
| 50 | -0.61 | 5.03 | -7.26 | 4.26 |

Table 4: Reference data for 50 species in the simulation study.
